# REP-X: An Evolution-guided Strategy for the Rational Design of Cysteine-less Protein Variants

**DOI:** 10.1101/797969

**Authors:** Kevin Dalton, Tom Lopez, Vijay Pande, Judith Frydman

**Affiliations:** Biophysics Program, Stanford University, Stanford, California, USA; Department of Biology, Stanford University, Stanford, California, USA; Department of Chemistry, Stanford University, Stanford, California, USA

## Abstract

Site-specific labeling of proteins is often a prerequisite for biophysical and biochemical characterization. Chemical modification of a unique cysteine residue is among the most facile methods for site-specific labeling of proteins. However, many proteins have multiple reactive cysteines, which must be mutated to other residues to enable labeling of unique positions. This trial-and-error process often results in cysteine-free proteins with reduced activity or stability. Herein we describe a general methodology to rationally engineer cysteine-less proteins. Briefly, natural variation across orthologues is exploited to identify suitable cysteine replacements compatible with protein activity and stability. As a proof-of-concept, we recount the successful engineering of a cysteine-less mutant of the group II chaperonin from methanogenic archaeon *Methanococcus maripaludis*. A webapp, REP-X (Replacement at Endogenous Positions from eXtant sequences), which enables users to design their own cysteine-less protein variants, will make this rational approach widely available.

## Introduction

Modification of a unique cysteine residue with a maleimide or iodoacetamide conjugated probe is a widely used method of producing site-specifically labeled proteins. These two chemistries have a long history in the literature starting from the first description of the reaction between iodoacetamide and thiols in 1933^1,2^. The value of this reaction to protein biochemistry was reified shortly thereafter in the first published reports of iodoacetamide’s reaction with purified, native proteins ^3,4^. The promise of maleimide chemistry was confirmed in 1949 when the reaction between substituted maleimides and thiols was shown to proceed at room temperature in water on the minute timescale with a nearly stoichiometric yield^5^. Shortly thereafter, the reaction of N-ethylmaleimide with actin was reported constituting the first published study to demonstrate that substituted maleimides react specifically with purified proteins^6^. In the following decade, creative applications of maleimide chemistry began appearing. The first deployment of bis-maleimides as a crosslinking agent was published in 1956^7^. Four years later, the first chromophore conjugated maleimide was used as an indicator for cysteine residues in proteolysis fragments of serum albumin^8^. Because of their high reaction yields under mild conditions, early biochemists appreciated the potential of substituted iodacetamides and maleimides to label proteins natively.

Today, cysteine directed probes are ubiquitous in protein science. Common substituents include crosslinkers, fluorophores, affinity tags such as biotin or digoxigenin, nitroxide spin labels, and silanes for surface immobilization. Such thiol-reactive probes are used in a range of applications including bulk kinetic measurements, single molecule biophysics, surface plasmon resonance, nuclear magnetic resonance and electron spin resonance spectroscopy^9^. A growing number of thiol-reactive probes are commercially available. As of writing, a search of Thermo Fisher’s online catalog alone contains over 90 thiol-reactive labeling kits and at least 13 thiol-reactive crosslinkers. Site-specific labeling has benefited the study of many protein families. However, in order to label thiols, one must first generate single cysteine mutants of a protein of interest. Typically, cysteines are replaced by serines or alanines, in an empirical trial-and-error process that often yields modified cysteine-less mutants that have reduced activity or stability (Figure 1A). Hence, a rational method to produce active cysteine-less protein variants would be quite desirable to contemporary protein science.

**Figure 1:**
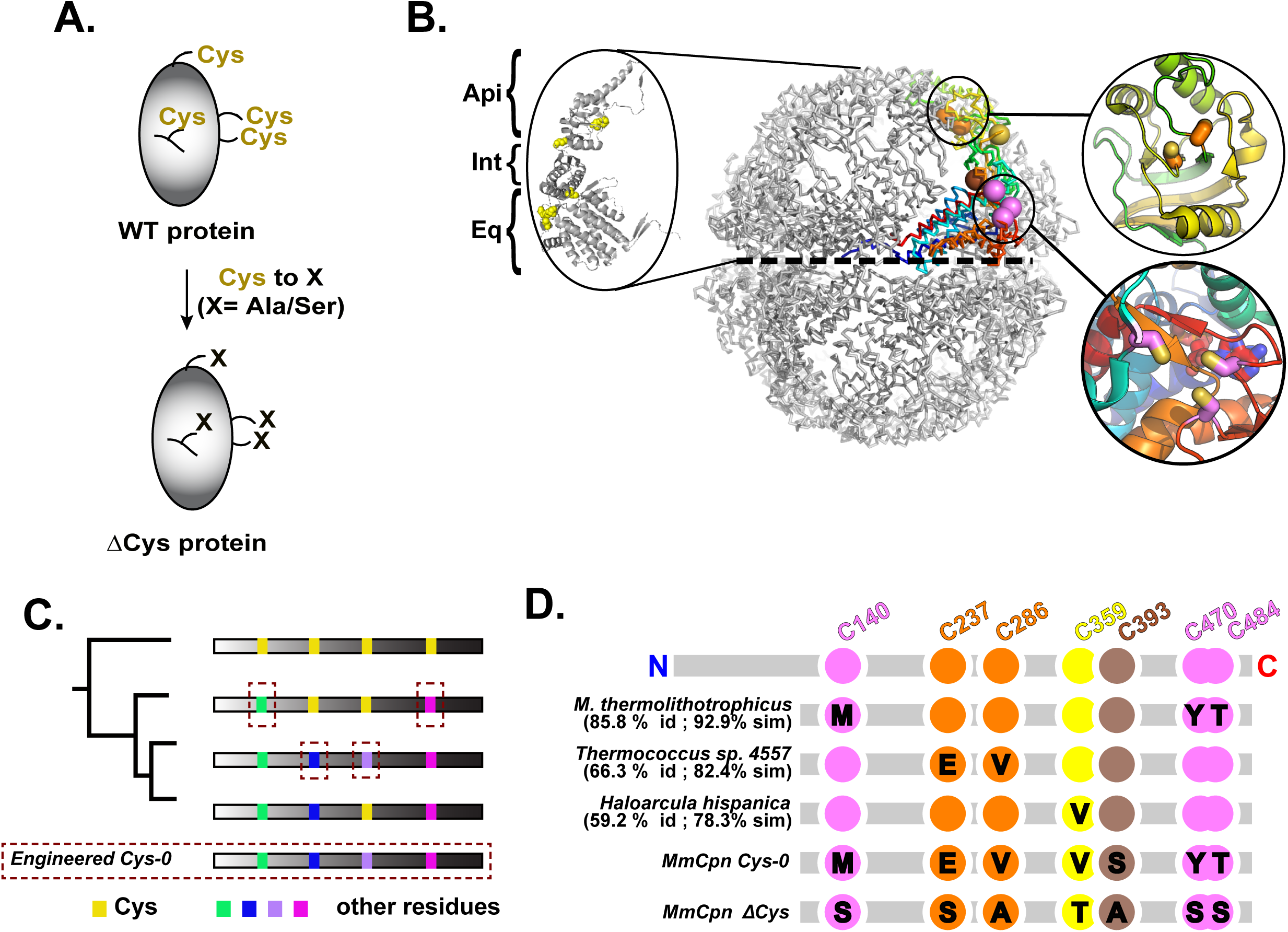
Design of Mm-Cys-0, a cysteine-less chaperonin variant. (A) Strategy for the design of Cysteine-less protein variants. (B) Molecular architecture of group II chaperonin Mm-Cpn. The crystal structure of the closed state of MmCpn-RLS mutant bound to ADP-AlFx (PDBID: 3RUW^24^. Highlighted are the equator of the complex (dashed line) and the three structural domains. A single subunit is shown in color with the seven cysteine residues rendered as spheres. Inset: cysteine clusters in the apical (top) and equatorial domains (bottom) (C) Design strategy of a cysteine-less MmCpn. Positions of seven Cys residues in Mm-Cpn WT are highlighted in yellow. (D) Extant sequences yielding the substitutions used to create the Cys-0 variant are shown along with the corresponding percentage identity and similarity between their sequence and MmCpn. The sequence of MmCpnCys-0 (this work) is shown as well as the sequence of MmCpnΔCys^16^.

Along these lines, we sought to generate a cysteine-less variant of a well studied group II chaperonin, MmCpn. Chaperonins are ATP-driven components of the cellular protein folding machinery and are essential in all free-living organisms. They comprise two structurally related chaperonin families generally called group I and group II chaperonins. Group I chaperonins are present in prokaryotes and the endosymbiotic organelles, chloroplasts and mitochondria^10^. Group I chaperonins are also, albeit comparatively rarely, found in archaeal genomes. The study of the archetypal group I chaperonin, GroEL from Escherichia coli, has long benefitted from the availability of a cysteine-less variant^11^ that enabled many mechanistic studies. Even for this successful example, the cysteine-less GroEL variants exhibit impaired activity compared to wild type^11^. Comparable studies have not been possible thus far in the group II chaperonins.

The group II chaperonins are found exclusively in the archaeal and eukaryotic cytosol^12^. Broadly, they are a family of ATP-driven hexadecameric chaperones with two stacked, eight membered rings per complex. These rings are related by a two-fold symmetry axis, which lies along the equator of the complex (Figure 1B, dashed line). Orthogonal to the two-fold axis is the eight-fold axis, which defines the intra-ring symmetry of the complex. The eukaryotic group II chaperonin, termed TRiC or CCT comprises eight parologous subunits of which four bind ATP. By contrast, the archaeal group IIs are prototypically α_8_β_8_ complexes. Other configurations have been reported including group IIs with 9-fold symmetry^13,14^. Here we will discuss the homohexadecameric group II chaperonin from *Methanococcus maripaludis*, MmCpn, which has proven itself to be a very useful model group II chaperonin.

All group II chaperonins share a conserved subunit architecture. The equatorial domain contains the nucleotide binding pocket. The remaining two structural domains of the monomer are the intermediate and apical domains arranged in ascending order from the complex equator (Figure 1B). The ring interface of MmCpn is formed by the equatorial domain of the chaperonin. The catalytic residue for ATP hydrolysis, D386 in MmCpn, is located in the intermediate domain. The apical domain binds misfolded client proteins. ATP hydrolysis leads to the release of client from the apical domains into the central cavity of the chaperonin. At this point in time, the apical domains close over the client forming an 8-fold symmetric iris and shielding the client from the bulk cytosol such that it my fold in isolation.

MmCpn monomers are 543 residues long and contain seven cysteines distributed throughout the primary sequence in all three structural domains of the enzyme. Cys-140, 470, & 484 are located in the equatorial domain near the ATP binding pocket. There are two cysteines in the intermediate domain, Cys-359 & 393. Finally, the apical domain contains a pair of cysteines, residues C237 & 286. The three equatorial cysteines are in steric contact with one another (Figure 1B bottom inset). Additionally the two apical cysteines are also in close proximity (Figure 1B top inset).

Previous efforts demonstrated that a cysteine-less mutant of MmCpn could be produced from a combination of serine, alanine and threonine point mutations^15,16^. This MmCpn mutant, termed ΔCys (MmCpnΔCys), assembles correctly and can bind unfolded proteins. It can also close its lid upon the addition of ADP-AlFx, which acts as a transition state analog for the ATP hydrolysis reaction. However, MmCpnΔCys is very poorly expressed in *E. coli*, and thus unsuitable for applications requiring milligram quantities of protein. To engineer a stable and highly expressed cysteine-less MmCpn variant for structural and biophysical work, we sought an alternative strategy to identify position-specific cysteine substitutions. We developed a rational approach using natural variation among extant ortholog sequences, which yielded a well expressed chaperonin with wild-type client folding activity. This effort provides proof-of-concept for our strategy to engineer amino acid replacements in proteins.

## Results

### A rational strategy to design a cysteine-less chaperonin variant

In order to design a functional, well-expressed cysteine-less chaperonin mutant we sought to integrate available structural information about MmCpn with phylogeny, using extant sequence information for homologous archaeal chaperonins. We reasoned that amino acid substitutions derived from close MmCpn homologs would be well tolerated by this chaperonin. To this end we amassed 2981 group II chaperonin sequences from the NCBI non-redundant protein sequence database using BLAST^17^. Each of these extant chaperonin sequences was aligned to the WT MmCpn sequence using the program Water from the EMBOSS Suite^18^. Water implements the Smith-Waterman sequence alignment algorithm which guarantees optimal pairwise local sequence alignment given an affine gap penalty^19^. To better interpret the information from the alignments, the sequences in the database were then rank ordered by their percentage identity with MmCpn.

The overall strategy of our approach to identify compatible amino acid replacements for each cysteine is outlined in Figure 1C. We carried out alignments to query the chaperonin sequence database for the most closely related sequence with a substitution at each cysteine position. These analyses informed the choice of residues to substitute to create a cysteine-less functional variant. One challenge in the implementation of this strategy was presented by sets of cysteines in steric contact, which in Mm Cpn are present in the equatorial and apical domains. Each of these sets consisted of a cluster of several cysteines in steric contact in the MmCpn structure. We were concerned that mutation of one of these residues would require compensatory mutations in the other(s) that maintained the structural integrity of their contact. Accordingly, when the database was queried for the equatorial cysteines (Figure 1B, D Magenta), we restricted hits to sequences with substitutions at all three positions. Likewise, for the two cysteines in the apical domain (Figure 1 B, D orange), we restricted hits to chaperonin sequences with both residues substituted.

Searching our database for non-cysteine variants of the equatorial cysteine positions C140, 470 & 484 returned a sequence from *Methanococcus thermolithotrophicus* with overall 85.8% identity to MmCpn but with non-cysteine residues at all three cysteine positions (Fig 1D, Magenta). Notably, this was the most closely related sequence in the database with non-cysteine variants at any of the three equatorial cysteine residues, suggesting that a mutation at any one of these positions necessitates a compensatory mutation at the other sites. The identities of the corresponding mutations in MmCpn are C140M, C470Y and C484T (Figure 1D). Similarly, searching for coincident non-cysteine variants of the two apical Mm-Cpn cysteines yielded a sequence containing the substitutions C237E and C286V from *Thermococcus sp. 4557* which is 66.3% identical to MmCpn (Figure 1D).

The remaining two cysteines are in the intermediate domain. Cys-359 was found to be mutated to valine in *Haloarcula hispanica* which is 59.2% identical to MmCpn. This was the most closely related organism with any substitution at that location. Finally, Cys-393 exhibited several amino acid substitutions in closely related sequences (Figure S1). As such, we chose to incorporate serine at this position because it is both present in extant sequences and is isosteric with cysteine. Notably, MmCpnΔCys incorporates an alanine at this position, which is also well represented in the database of homologs. All the above substitutions were introduced into MmCpn to generate a new cysteine-less variant herein termed ***Cys0*** *(MmCpn-Cys0)*.

### The cysteine-less MmCpn-Cys0 is well expressed and supports ATP-driven cycling

While the previously derived MmCpnΔCys could assembled correctly, it did not express well in *Escherichia coli*.This is illustrated by experiments where wild type (WT) and ΔCys MmCpn were expressed from pET21a plasmids in *E. coli* BL-21. Following induction with IPTG, expression was monitored by SDS-PAGE of total lysates (Figure 2A). WT MmCpn was highly expressed in soluble form and clearly visualized by Sypro Ruby staining, while the previously described CpnΔCys mutant protein was barely perceptible in the lysate. Importantly, the same experiment demonstrated that the rationally designed MmCpn-Cys0 variant was produced at much higher expression levels, comparable to WT (Figure 2A). In a further test of assembly competence, we examined whether the Cpn variants remained soluble after treatment of the lysate with 55% ammonium sulfate (AS), as observed for the WT MmCpn (Figure 2B). Indeed, both WT and Cys-0 variants, but not the ΔCys variant, were very soluble in 55% ammonium sulfate (Figure 2B). Accordingly Cpn Cys-0 could be purified with the MmCpn purification protocol established for WT.

**Figure 2:**
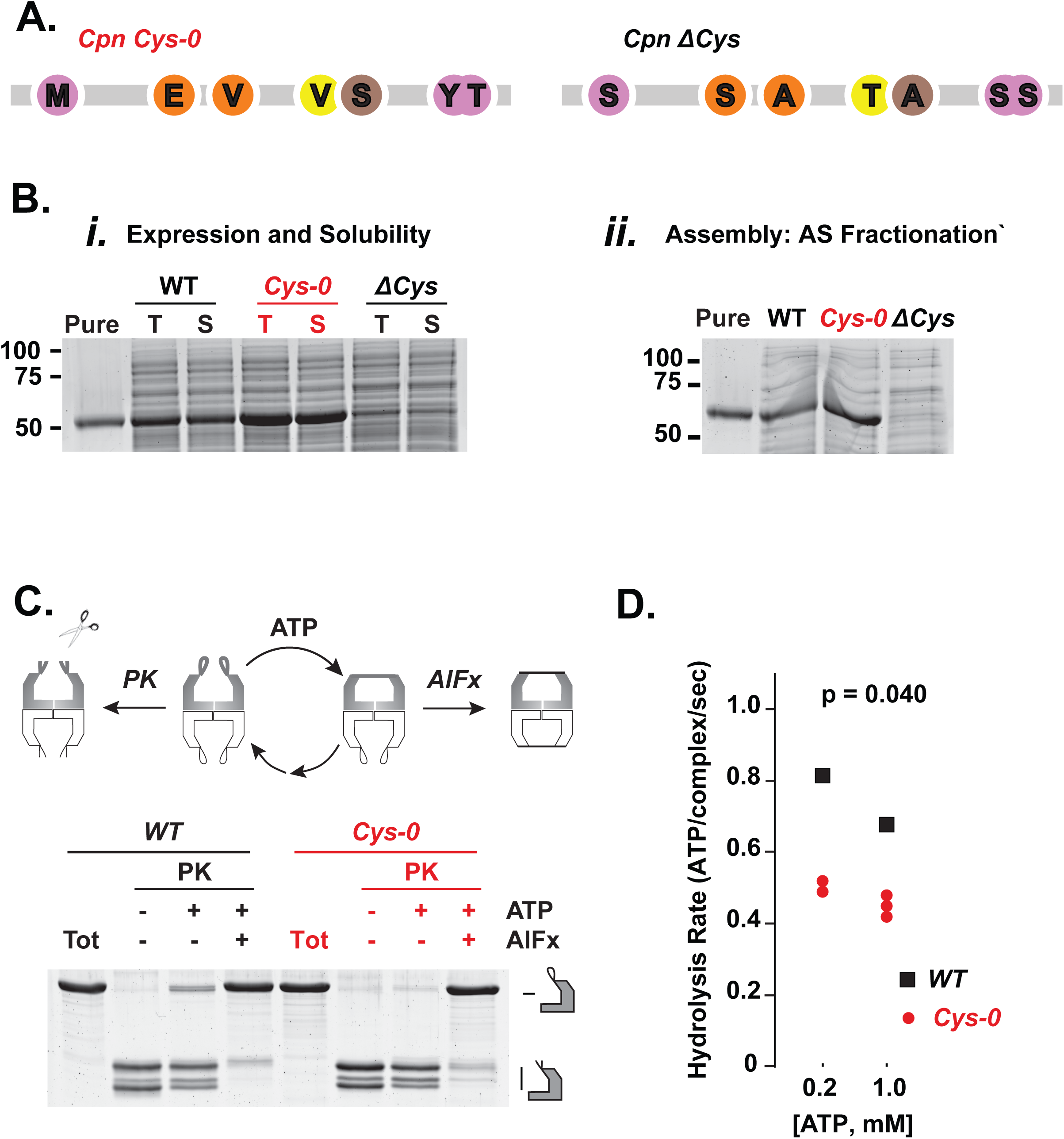
MmCpnCys-0 is well expressed and has ATPase activity. (A) Cys replacements in MmCpnCys-0 and MmCpnΔCys. (B) Test expression of chaperonin variants. Lane 1, 1ug of purified WT MmCpn. Other lanes contain 2 ul of each sample. T indicates total lysate while S indicates soluble proteins following centrifugation at 22,000 × g for 30 minutes. (C) Ammonium sulfate fractionation as a test of complex assembly. SDS-PAGE of 2 ul of test expressions following a 55% ammonium sulfate cut. (D) Proteinase K protection assay for purified chaperonin mutants. All gels stained with Sypro Ruby (E) ATPase assay for MmCpnCys-0. Mean WT ATPase rates indicated by horizontal lines.

We next examined whether MmCpn Cys-0 has ATPase activity and can undergo the ATP-driven conformational change typical of Group II chaperonins. ATP hydrolysis promotes a conformational change from an open, client binding conformation to a closed configuration, characterized by formation of a built-in lid over the central chamber of the complex (Figure 2C). To this end, we exploited the observation that, in the open state, the lid segments are sensitive to non-specific proteolytic digestion by proteinase K, while in the closed state these lid segments are strongly protease-protected (Figure 2C). Inclusion of AlFx, a transition state analogue, in the ATP hydrolysis reaction stabilizes the closed conformation, rendering the chaperonin resistant to Proteinase K. This assay demonstrated that incubation with ATP and AlFx yielded similar levels of protection to both WT and Cys-0 chaperonins, indicating the Cpn Cys-0 mutant can fully attain the closed conformation in response to ATP (Figure 2C). As described below (Figure 3A), native agarose gel electrophoresis confirmed this conclusion and provided additional support for the ATP-induced formation of the closed Cpn conformation (Figure 3A bottom panel).

**Figure 3:**
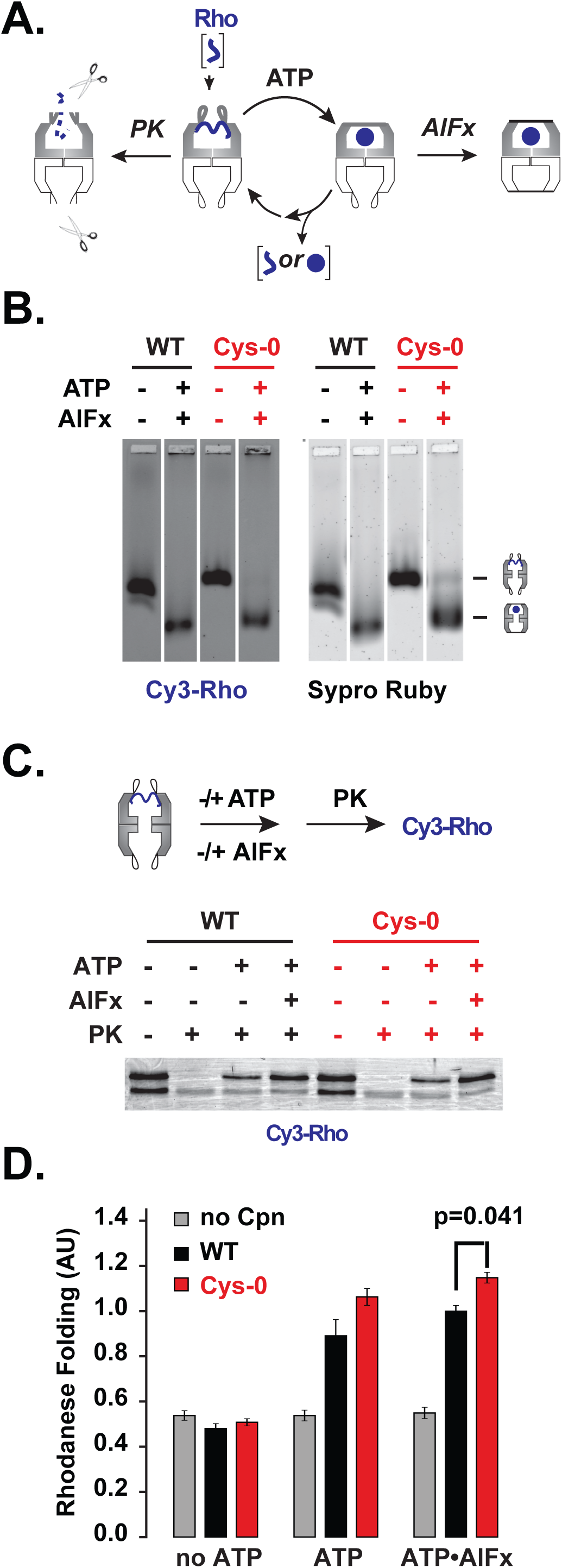
MmCpnCys-0 can effectively mediate substrate folding. (A) Scheme of chaperonin folding cycle (B) Non-denaturing agarose electrophoresis to monitor binding MmCpn and the substrate protein rhodanese. Cy3-maleimide labeled rhodanese co-migration with MmCpn in different nucleotide states (top). Total protein stained with Sypro Ruby (bottom). (C) Protease protection of Cy3-rhodanese by MmCpn in different nucleotide states. (D) Rhodanese folding of WT and Cys-0 MmCpn. The y-axis represents the amount of cyanide converted to thiocyanate by rhodanese in 10 minutes relative to the mean amount converted by the wild type sample with ATP-AlFx.

Without the addition of AlFx, the chaperonin will freely hydrolyze ATP and cycle between the open and closed conformation in the process. The amount of proteinase K protection observed in this state thereby reports on the fraction of its nucleotide cycle the chaperonin spends in the open, proteinase sensitive state. Under cycling conditions, the Cys-0 variant was less proteinase-protected than WT, suggesting that MmCpnCys-0 spends more time in the open state during its nucleotide cycle (Figure 2C). This observation is consistent with a decreased rate of chamber closure or an increase in the rate of chamber re-opening. To evaluate the likelihood of each explanation, we measured the ATP hydrolysis rate of MmCpn Cys-0 compared to WT. For all three ATP concentrations tested, the Cys-0 variant hydrolyzed ATP at comparable, albeit slightly lower rates that WT Cpn (Figure 2D). This result strongly suggests that the decreased level of proteinase protection seen in cycling Cys-0 is due to a decreased rate of closure and not faster re-opening which would manifest and increased ATPase rate.

Of note, both WT and Cys-0 chaperonins exhibit the previously described inhibition of ATP hydrolysis rates at high ATP concentrations (compare 200 and 1000 micromolar ATP in Figure 2D). This indicated that MmCpnCys-0 preserves the allosteric regulation of ATP hydrolysis characteristic of group II chaperonins.

### MmCpn Cys-0 is fully functional in ATP-dependent substrate folding

Next, we examined the ability of MmCpn Cys-0 to mediate protein folding. Bovine rhodanese, responsible for detoxifying cyanide ions in vivo, is a well characterized model substrate for MmCpn. Once denatured, rhodanese cannot fold without chaperone assistance, forming insoluble aggregates instead. However, denatured Rhodanese diluted into buffer containing WT MmCpn binds to the chaperonin, forming a characteristic high molecular weight complex (Figure 3A). Upon addition of ATP or ATP-AlFx to induce lid closure, Cpn encapsulates rhodanese within its central chamber. Rhodanese folds to the native active state within the central chaperonin chamber, and is released upon reopening of the chamber following ADP and phosphate release (Figure 3A) ^20^. With ATP-AlFx lid reopening is blocked, and the encapsulated substrate remains locked within the closed chaperonin chamber.

To compare the abilities of WT and Cys-0 Mm-Cpn to fold Rhodanese in an ATP-dependent manner, we dissected substrate engagement at different steps of the ATP-driven Cpn folding cycle (Figure 3B-D). We first examined substrate binding using denatured cy3-labeled rhodanese. Cpn binding was monitored by comigration with Rhodanese on non-denaturing gel electrophoresis. Protein staining visualized the slowly migrating Cpn (Sypro Ruby, Figure 3B) while fluorescence imaging visualized Cy3-rhodanese (Figure 3B). Both variants formed the characteristic Rhodanese-Cpn complex in the absence of ATP, with similar efficiency. Upon incubation with ATP-AlFx, both WT and Cys-0 similarly changed their mobility to the faster migrating lid-closed state. Since WT and Cys-0 comparably retained the bound substrate in this analysis, we concluded MmCpn Cys-0 effectively encapsulated substrate upon closure.

The ability of Cys-0 to encapsulate the bound Rhodanese was tested directly using a protease sensitivity assay. A rhodanese-Cpn complex was treated with Proteinase K following incubation in the absence or presence of ATP or ATP-AlFx (Figure 3C). In the open state, the Cpn-bound non-native protein is unstructured and extremely protease sensitive. However, upon ATP-induced lid closure, the encapsulation of the substrate within the chamber results in full protection from proteolysis. As expected, rhodanese bound to either WT or Cys-0 Cpn was protease sensitive. Incubation of either WT or Cys-0 with ATP-AlFx or with ATP led to full protection of the bound substrate from proteinase K digestion. This indicates that ATP leads to efficient encapsulation of the bound substrate within the closed central chamber both for WT Cpn and for the Cys-0 variant (Figure 3C).

Finally, we compared the ATP-dependent production of folded Rhodanase when bound by MmCpn Cys-0 versus MmCpn WT (Figure 3D). Rhodanese folding results in active enzyme that can be measured using a colorimetric assay. The rhodanese-Cpn complex was incubated in the absence or presence of ATP or ATP-AlFx, and the extent of rhodanese folding was measured using an endpoint assay. Importantly, the cysteine-less MmCpn-Cys-0 promoted rhodanese folding as effectively as WT Cpn both under cycling conditions,i.e. in the presence of ATP, or when the complex is locked in the closed state with ATP-AlFx. Together, these experiments indicate that our approach enabled the design of a well folded, assembled and active cysteine-less chaperonin complex.

## Discussion

We here describe a rational bioinformatic approach to engineer active cysteine-less variants of proteins using an analysis of extant sequences. Our premise is to exploit natural variation to identify amino acid substitutions compatible with a protein’s structure and activity. Unless cysteines are essential for protein catalysis or to generate a specific structural or functional motif, such as a Zn-finger, it is likely that examination of orthologues of a protein of interest will identify suitable cysteine replacements compatible with its structure and activity. We documented the success of our strategy by engineering a cysteine-less variant of the group II chaperonin MmCpn. To our great surprise, this technique allowed us to define seven simultaneous point mutants that together could replace the cysteines in the protein while retaining a functional, well expressed complex. The unintuitive nature of the point mutations selected by this approach is worth emphasizing.

The common practice to eliminate cysteines by mutation to alanine or the isosteric amino acid serine often fails to yield active or stable protein variants. This is illustrated by analysis of the previously obtained cysteine-less Cpn mutant ΔCys, which is less stable and less well expressed, indicating this approach is not the best choice for the chaperonin system.

Furthermore, to enumerate and experimentally explore all possible, 20^7 (1.28 trillion), amino acid combinations is too onerous to be feasible. A remarkable feature of our approach is its ability to identify spatially or structurally linked replacements. For instance, this allowed us to replace cysteine clusters that are in steric contact in the folded protein. By identifying sets of non-cysteine residues that maintained the interaction in orthologous proteins, our approach permitted the replacement of the entire cysteine cluster without loss of protein integrity or activity.

To make this engineering approach available to other investigators we developed a software tool which we named Replacement at Endogenous Positions from eXtant sequences (REP-X). The current (alpha) version of the software is available at (http://github.com/kmdalton/rep-x). The basic design of the application is summarized in Figure 4. We predict this software will enable other protein scientists to rapidly and rationally design cysteine-less protein variants, opening the way to introducing cysteines at specific positions for site specific labeling.

**Figure 4:**
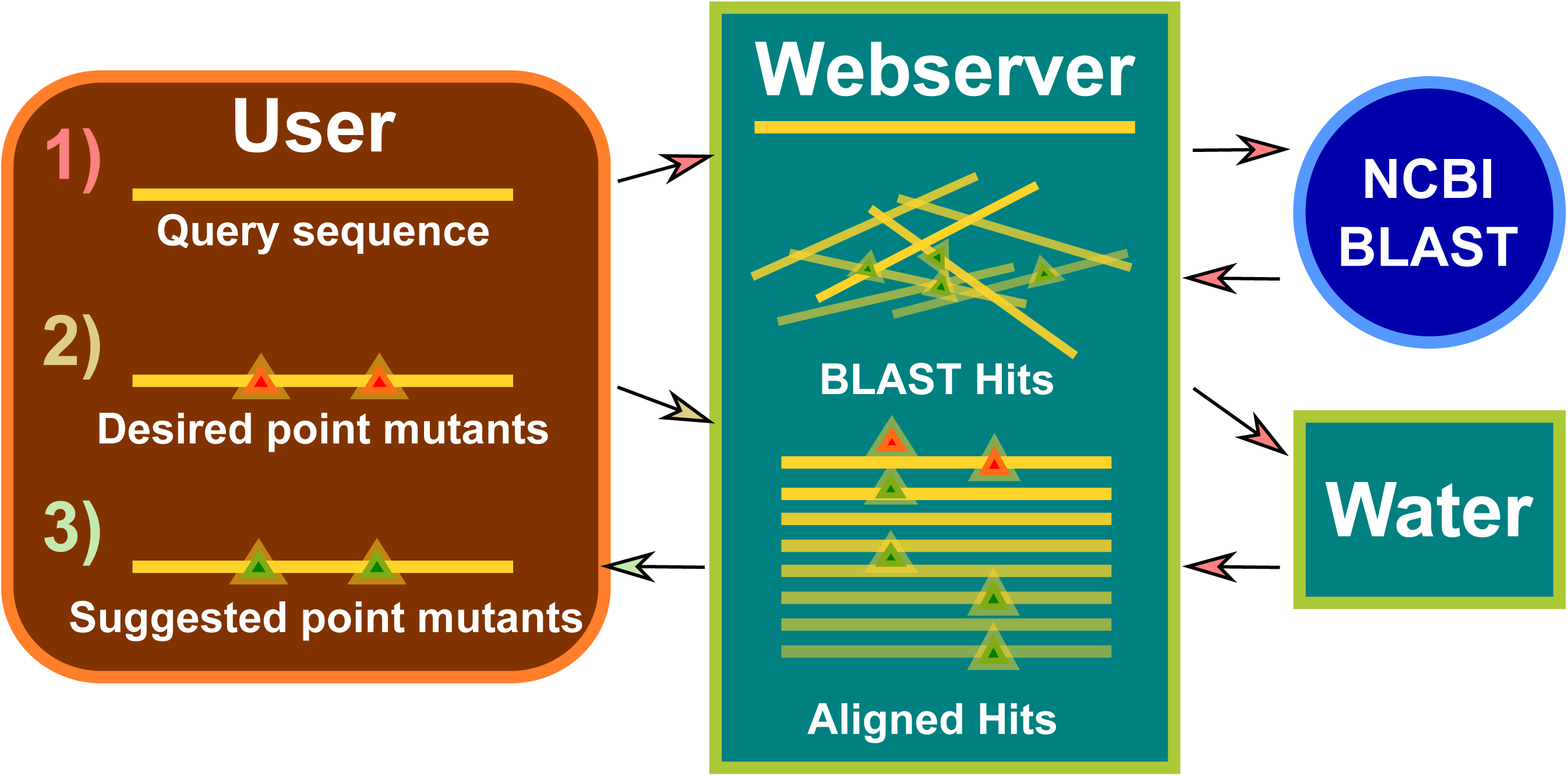
Design of the Rep-X Webserver. 1) Query sequences are submitted to the webserver by users. BLAST^17^ hits are retrieved by the NCBI server and aligned to the query sequence using Water from the EMBOSS Suite^18^. 2) Point mutants are requested by the user, and 3) point mutants are suggested by the webserver.

An important consideration for our approach, is that there is no intrinsic limitation on the amino acid type being replaced. REP-X can equally aid in the design of single tryptophan mutants for intrinsic fluorescence studies, be used to generate single lysine variants for site specific labeling by N-hydroxysuccinimide conjugates or to generate specific single amino acid variants for position specific labelling with stable isotopes for NMR experiments. Our results suggest that this technique can be particularly powerful in systems for which structural data are available in order to identify cysteines which are in close proximity.

## Materials & Methods

### MmCpn Expression Constructs

MmCpn WT and MmCpnΔCys plasmids derive from previous work. Briefly, the sequence of the chaperonin was cloned into the Nde1 and BamHI of pET21a+ (EMD Chemicals) yielding pET21MmCpnWT^21^. MmCpnΔCys was obtained from D. Hoersch and T. Kortemme^16^. The C-terminal polyhistidine tag of MmCpnΔCys was removed by the introduction of a stop codon using the QuickChange II site-directed mutagenesis kit (Agilent Technologies). Codon optimized MmCpn Cys-0 in the pJ414 vector was obtained from DNA2.0.

### Buffer Composition

Three standard chaperonin buffers were used in this work. MQA: 20 mM HEPES-KOH (pH 7.4), 50 mM NaCl, 5 mM MgCl_2_, 0.1 mM EDTA, 10% (v/v) glycerol. MQB: 20 mM HEPES-KOH (pH 7.4), 1 M NaCl, 5 mM MgCl_2_, 0.1 mM EDTA, 10% (v/v) glycerol. ATPase buffer: 20 mM Tris-HCl (pH 7.5), 100 mM KCl, 5 mM MgCl_2_, 10% (v/v) glycerol). Buffers were supplemented with 1mM DTT and 0.5 mM PMSF during lysis.

### Expression tests of chaperonin mutants

E. coli BL-21 pRosettas containing the three expression vectors were cultured in terrific broth with 100 ug/ml ampicillin and 30 ug/ml chloramphenicol overnight then diluted 1:1000 into 50 ml of fresh media. These cultures were grown at 37 C with 250 rpm shaking until reaching an OD600 between 0.9 and 1.0. The cultures were induced with 1mM IPTG and transferred to a 16 C incubator overnight. The following morning the cells were pelleted by centrifugation at 5,000 × g for 10 minutes and flash frozen in liquid nitrogen.

Frozen cell pellets were resuspended in 10 ml of MQA buffer and lysed in an Emulsiflex-B15 (Avestin). Lysates were cleared by centrifugation at 22,000 × g for 30 minutes. Ammonium sulfate was added to the cleared lysate to a final concentration of 55% saturation. After a 15 minute incubation at 4 C, the insoluble fraction was removed by centrifugation at 22,000 × g for 30 minutes. 2 ul of the lysed, cleared, and ammonium sulfate soluble fractions were loaded onto 15% Tris-glycine SDS Polyacrylamide gels and electrophoresed at 25 mA for 1 hour. Gels were stained with Sypro Ruby (ThermoFisher) and imaged on Typhoon scanner (GE).

### Chaperonin purification

Expression, lysis, and ammonium sulfate cuts were performed as described in the previous section excepting that the culture volumes were increased to 1 L. Anion exchange and heparin affinity chromatography were performed as described previously^21^. After heparin chromatography, MmCpn solutions were concentrated in 15 ml centrifugal concentrators with a 100 kDa nominal molecular weight cutoff, buffer exchanged at least 3 times into MQA, concentrated to between 80 and 150 mg/ml, flash frozen in liquid nitrogen and stored at −80 C.

### ATPase assay

A previously described enzyme coupled assay ^22^ was used to measure the ATPase activity of MmCpnCys-0. 180 uL of ATPase buffer containing 10 mM phosphoenolpyruvate, 1.5 mM NADH, 5 units of pyruvate kinase (rabbit muscle type VII buffered aqueous glycerol solution, Sigma-Aldrich), 4.6 units of l-lactic dehydrogenase (bovine heart type XVII buffered aqueous glycerol solution, Sigma-Aldrich), and 0.25 M MmCpn was warmed to 37C on a heat block. This was mixed rapidly with the indicated concentration of ATP and pipetted into a quartz cuvette which was pre-equilibrated at 37C in a Hewlett Packard 8453 spectrophotometer. Conversion of NADH to NAD+ was monitored by a decrease in absorbance at 340 nm for 120 seconds. The linear region of the timecourse was fit and the slope used to calculate the rate of ATP hydrolysis by MmCpn.

### Proteinase K assays

Assays were performed as described previously ^21^ except that the chaperonin nucleotide reaction incubation time was extended to 60 minutes at 37 C. Briefly, ATPase buffer containing 0.25 uM MmCpn was pipetted onto the appropriate ATP conditions (no ATP, 1 mM ATP, 1 mM ATP + 1 mM AlNO3 + 6 mM NaF), and incubated at 37 C for 1 hour. Subsequently, 400 ng of proteinase K was added and the chaperonin was digested at room temperature for 10 minutes. The reaction was quenched by the addition of 0.05 M phenylmethanesulfonyl fluoride. The samples were electrophoresed on a 15% tris-glycine SDS-PAGE gel and stained with Sypro Ruby (Thermo Fisher).

Cy-3 rhodanese protection was performed just as the standard proteinase K protection assay except that 0.25 uM chaperonin was first added to 1 uM final concentrated rhodanese dissolved in 6mM guanidium buffered with 100 mM HEPES pH 7.4 for 10 minutes on ice and subsequently centrifuged for 10 minutes at top speed in a microcentrifuge. This sample was applied to the nucleotide conditions, proteinase, and quenched with PMSF as described above.

### Rhodanese folding assay

Rhodanese folding was performed as described previously ^20^. Briefly, 100uM denatured rhodanese in 100 mM HEPES pH 7.4 with 6M guanidinium chloride and 5 mM DTT was diluted 1:100 into ATPase buffer containing 0.25 uM MmCpn. Samples were incubated on ice for 10 minutes followed by a 10 minute spin at top speed in a 4C refrigerated microcentrifuge to pellet aggregated rhodanese. The samples were pipetted onto nucleotide conditions containing no ATP, 5 mM ATP, or 5 mM ATP + 1mM AlNO3 + 6 mM NaF and incubated for 60 minutes at 37 C. Rhodanese activity was then assayed colorimetrically as described previously ^23^. A single 10 minute time point was used. 0.5 M EDTA was substituted for 0.4 M CDTA.

### Non-denaturing agarose gel electrophoresis

Samples were prepared as in the proteinase protection assays, but rather than being digested, they were loaded onto a 1% agarose gel with 1 mM MgCl_222_ and 80 mM MOPS pH 7.4, and electrophoresed for 1 hour at 100 volts and optionally stained with Sypro Ruby (Thermo Fisher) before imaging staining on a Typhoon Scanner (GE).

### Rhodanese Purification

10 mg of bovine rhodanese (Sigma) was resuspended in 4 ml of buffer A (50 mM acetate pH 5.0 with 20 mM sodium thiosulfate). Undissolved material was removed by centrifugation at top speed in a microcentrifuge. The supernatant was applied to a Mono-S chromatography column and eluted over a linear gradient into buffer B (50 mM acetate, pH 5.0, 20 mM sodium thiosulfate, and 500 mM NaCl). The fraction containing the most rhodanese was further purified by size exclusion chromatography on an Superdex-75 column pre-equilibrated with buffer A. Resulting fractions were dehydrated in a speedvac and resuspended to a final concentration of 100 uM in 6M guanidinium with 100 mM HEPES at pH 7.4.

### Cy3-Rhodanese Labeling

After Mono-S chromatography, 400 ul of rhodanese solution was reserved from the purification. This solution was added onto a solution of Cy3-Maleimide dissolved in 20 ul of DMSO and incubated at room temperature for 2.5 hours. The labeling reaction was quenched with 10 ul of 1M DTT. This solution was then purified by size exclusion and denatured at described above.

## Acknowledgements

We thank D. Hoersch and T. Kortemme for the plasmid encoding MmCpnΔCys and members of the Frydman lab for useful discussions. This work was funded by NIH (GM074074).

## Author contributions

KD wrote the program and carried out most experiments in collaboration with TL. VP and JF provided guidance, JF directed the project. KD and JF wrote and all authors edited the MS.

## Competing interests

None

## Data availability

REP-X available at http://github.com/kmdalton/rep-x

## Supplemental Figures

**Figure S1:**
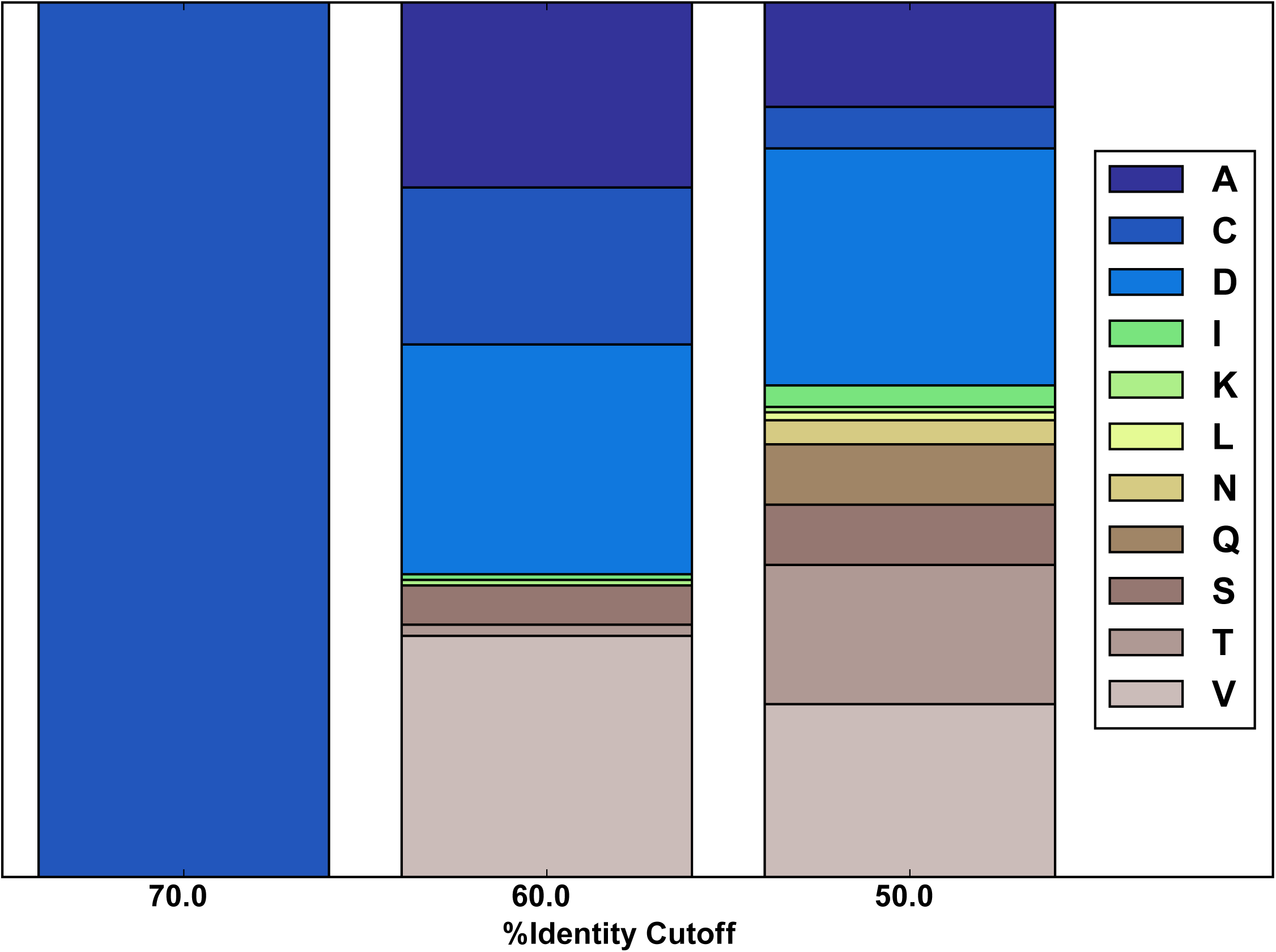
Distribution of Residues at Position 393 in Cpn orthologues. The observed distribution of amino acids at position 393 for protein sequences in the MmCpn homolog database. Distributions are shown for homologs with at least 70, 60, or 50 % sequence identity with MmCpn.

## Notes

http://github.com/kmdalton/rep-x

